# LINC01503-MP is a Mitochondrial Microprotein That Promotes Cell Proliferation and Oxidative Metabolism

**DOI:** 10.1101/2025.03.19.644091

**Authors:** Nikita Dewani, Jorge Ruiz-Orera, Oliver Popp, Ning Liang, Masanari Sugarawa, Jana F. Schulz, Franziska Witte, Clara Sandmann, Takahiro Tsuji, Susanne Blachut, Takaharu Katagiri, Ivanela Kondova, Sae Owada, Shinji Yoshii, Hiroshi Kataoka, Andreas Kurtz, Hiroshi Nakase, Sebastiaan van Heesch, Philipp Mertins, Norbert Hübner, Masatoshi Kanda

## Abstract

Long non-coding RNAs (lncRNAs) are well-established as key regulators of gene expression. However, emerging evidence reveals that some lncRNAs can also encode functional microproteins. In this study, we report the identification of an evolutionarily young microprotein encoded by *LINC01503*, expressed across several human tissues. This microprotein, designated as LINC01503-MP, localises to the mitochondria and exerts a proliferative effect on HCT116 colorectal cancer (CRC) cells. Functional studies reveal that LINC01503-MP regulates mitochondrial oxygen consumption rate, linking its activity to enhanced metabolic functions and cell proliferation. Interactome analyses identified multiple mitochondrial metabolism-related proteins as potential interaction partners. Our findings show that LINC01503-MP plays a role in the proliferative phenotype associated with *LINC01503* upregulation in CRC, suggesting the functional significance of evolutionarily young, lncRNA-derived microproteins in cancer progression.

## INTRODUCTION

Long non-coding RNAs (lncRNAs) represent a class of RNA transcripts exceeding 200 nucleotides in length that do not contain annotated protein-coding sequences (Mercer, Dinger and Mattick, 2009; Ponting, Oliver and Reik, 2009; Fu *et al*., 2016). LncRNAs are known to exert pivotal regulatory functions in gene expression (Cabili *et al*., 2011; Derrien *et al*., 2012; Volders *et al*., 2019; Senís *et al*., 2021; Statello *et al*., 2021; Gupta and Challagundla, 2022; Mattick *et al*., 2023). LncRNAs are involved in diverse regulatory mechanisms, acting as molecular scaffolds, guides, or decoys, influencing chromatin-modifying complexes and modulating transcription factor activity (Mattick *et al*., 2023), and interacting with RNA-binding proteins and transcription factors, thereby regulating mRNA stability, splicing, and other post-transcriptional processes (Ponting, Oliver and Reik, 2009). LncRNAs have garnered significant attention in cancer research due to their ability to regulate oncogenes or tumour suppressor genes. Dysregulation of lncRNA expression can drive tumour progression, invasion, and therapeutic resistance, highlighting their potential utility as diagnostic markers and therapeutic targets in cancer treatment (Gupta and Challagundla, 2022).

The discovery of translation from lncRNAs and the presence of unannotated non-canonical open reading frames (ncORFs) within these transcripts has challenged the traditional view that lncRNAs are strictly non-coding (Andrews and Rothnagel, 2014; Aspden *et al*., 2014; Ruiz-Orera *et al*., 2014; Ruiz-Orera, Villanueva-Cañas and Albà, 2020; Cai *et al*., 2021; van Heesch *et al*., 2019; Pan *et al*., 2022; Broeils *et al*., 2023; Xiao *et al*., 2024). Techniques such as ribosome profiling (Ribo-seq) (Ingolia *et al*., 2009) and mass spectrometry (Mohsen, Martel and Slavoff, 2023; Deutsch *et al*., 2024) have revealed that many lncRNAs harbour small ncORFs that can be translated into stable functional microproteins that are typically less than 100 amino acids (AA) in length (van Heesch *et al*., 2019; Pan *et al*., 2022; Broeils *et al*., 2023). For instance, *LINC00998*, a previously annotated lncRNA, encodes the microprotein SMIM30. This microprotein is shown to be a key regulator of hepatoma cell proliferation (Yang *et al*., 2023). Silencing SMIM30 reduces tumour growth and alters calcium signalling, while its overexpression promotes cell cycle progression by enhancing SERCA activity and lowering cytosolic calcium levels. Similarly, a microprotein encoded by *HOXB-AS3* has been shown to suppress colon cancer growth (Huang *et al*., 2017). The detection of such microproteins highlights the expanding complexity of the genome, suggesting that lncRNAs, beyond their regulatory non-coding roles, also contribute to the proteome through the generation of previously unrecognised small proteins. It also shows emerging evidence for the involvement of microproteins in various cancer mechanisms (Hofman, Prensner and van Heesch, 2024).

Additionally, large-scale screenings systematically identified biologically active ncORFs potentially encoding for microproteins in the human genome, many of which are annotated as lncRNAs. Clustered regularly interspaced short palindromic repeats (CRISPR)-based techniques have revealed hundreds of ncORFs shown to exert significant roles in cell survival. (Zheng *et al*., 2023; Delaidelli, Oliveira de Santis and Sorensen, 2024; Hofman *et al*., 2024). For example, one ncORF encoded by *GREP1* was found to translate a secreted protein highly expressed in breast cancer, with its knockout causing selective growth defects in breast cancer cell lines (Prensner *et al*., 2021). These findings underscore the potential of ncORFs encoded by lncRNAs as promising therapeutic targets or diagnostic biomarkers (Hofman, Prensner, and van Heesch, 2024). However, much remains unknown about these ncORFs, particularly regarding which specific ncORFs encode for stable and functional microproteins implicated in various cancers and diseases, as well as the molecular mechanisms underlying their associated phenotypes.

Recent studies have highlighted the emergence of evolutionarily young human microproteins across primate evolution that are integral to previously unrecognised biological pathways (Vakirlis *et al*., 2022; Broeils *et al*., 2023; Sandmann *et al*., 2023; Ruiz-Orera *et al*., 2024). Notably, many of these microproteins are encoded by ncORFs in lncRNAs and localise to the mitochondria, interacting with evolutionarily conserved proteins involved in essential cellular processes (van Heesch *et al*., 2019; S. Zhang *et al*., 2020; Sandmann *et al*., 2023). These facts suggest that these microproteins can significantly influence key cellular functions (van Heesch *et al*., 2019; Sandmann *et al*., 2023; Ruiz-Orera *et al*., 2024). This rapidly evolving field emphasises the importance of recently evolved lncRNA-derived microproteins in enhancing our understanding of protein diversity across evolution and their potential implications in health and disease.

In this study, we report a novel young ncORF-encoded microprotein derived from *LINC01503*, which we have named LINC01503-MP. Through a series of functional *in vitro* assays, we demonstrate the role of LINC01503-MP in promoting cell proliferation and its involvement in regulating mitochondrial oxygen consumption rate (OCR). We further investigated whether the LINC01503-MP microprotein is associated with cancer, particularly focusing on its protein expression in various tissues, including colorectal cancer (CRC). This work supports that the evolutionarily young microprotein derived from a previously annotated lncRNA may play significant roles in cancer progression.

## RESULTS

### Identification of a translated ncORF encoded by human *LINC01503*

To identify novel ncORFs potentially encoding for microproteins on lncRNAs, we retrieved four publicly available Ribo-seq datasets corresponding to healthy human tissues from four organs: kidney (n=6), heart left ventricle (LV) (n=15), liver (n=3) (van Heesch *et al*., 2019), and brain (n=3) (Z.-Y. Wang *et al*., 2020). These datasets were re-analysed by ORFquant (Calviello, Hirsekorn and Ohler, 2020) and PRICE (Erhard *et al*., 2018) (**Fig. 1a**, see **Methods**). We identified a total of 635, 637, 784, and 951 ncORFs on lncRNAs (lncORFs) in the kidney, heart, brain, and liver, respectively (**Fig. 1b**). These lncORFs have an average size of 50 amino acids (**Fig. 1c**), and only a small fraction of these lncORFs (3.17%, 72 out of 2,260) were identified across all four organs (**Fig. 1b**).

**Fig. 1.**
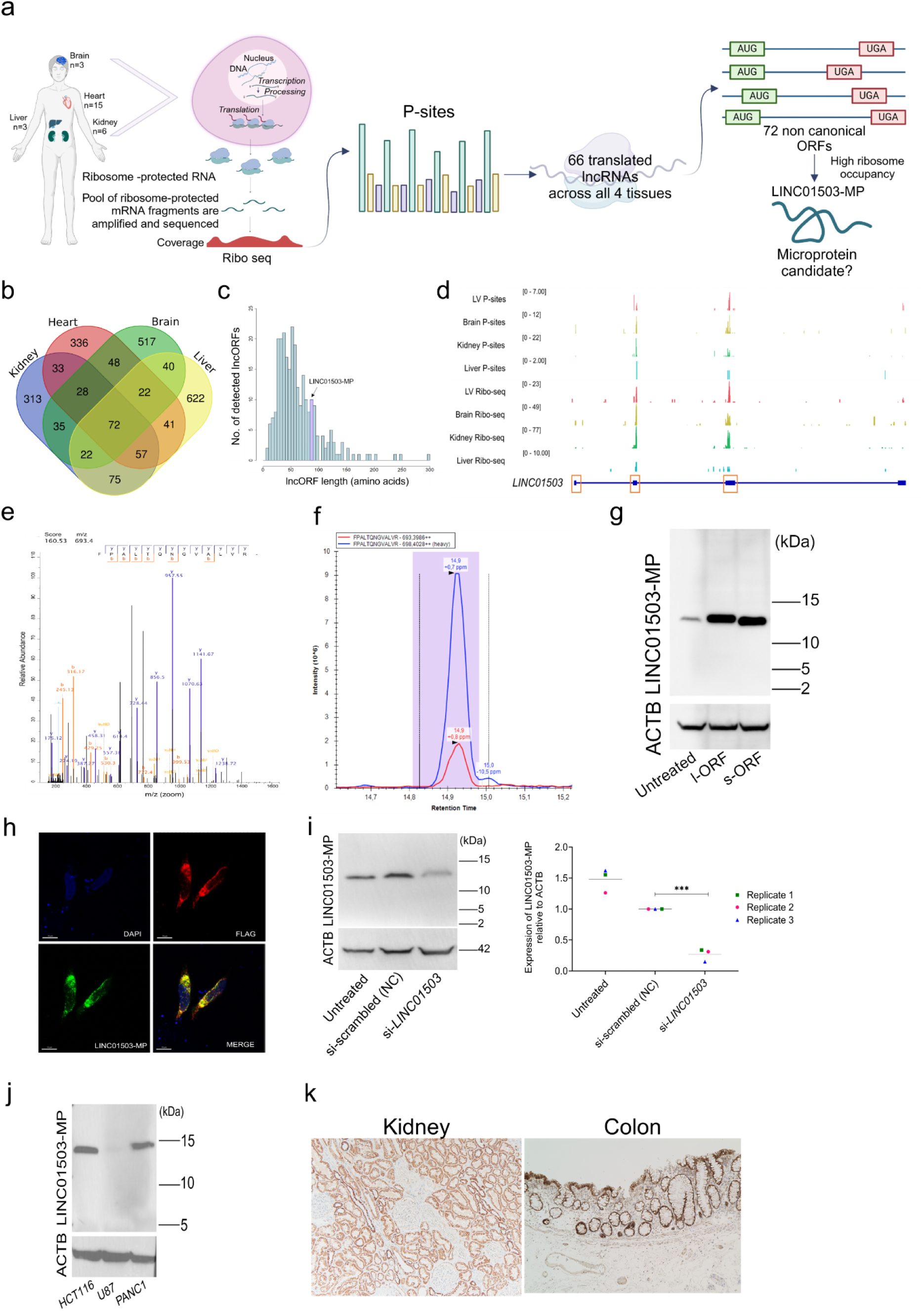
Candidate selection and detection of LINC01503-MP. a. Schematic overview of the microprotein candidate selection using Ribo-seq of human brain, heart, kidney and liver. b. Venn diagram of detected lncORFs in the four considered human organs; brain, heart, kidney and liver. c. Histogram depicting the number of lncORFs detected, highlighting the length of the long ncORF (l-ORF) encoded by *LINC01503*. d. Genome coverage tracks visualizing Ribo-seq reads and P-site positions on human *LINC01503* in the four considered tissues. The number inside of the square bracket indicates the range of counts. Orange boxes indicate the l-ORF encoded by *LINC01503*, spanning three exons. e. Identification of the unique peptide ‘FPALTQNGVALVR’ for LINC01503-MP by PRM in human kidney. f. Verification of the unique peptide (heavy) of LINC01503-MP in the human kidney by PRM mass spectrometry. g. Representative western blotting by anti-LINC01503-MP Ab and actin (ACTB) of HEK293T without (untreated) or with over-expressed s-ORF or l-ORF of LINC01503-MP. Similar results were obtained in 3 independent experiments. h. Representative immunofluorescence (IF) images of N-terminal 3xFLAG fused LINC01503-MP overexpressed HeLa cells by confocal microscopy: FLAG (red), anti-LINC01503-MP Ab (green) and DAPI (blue). Scale bar: 10 μm. Similar results were obtained in 3 independent experiments. i. Representative western blotting by anti-LINC01503-MP Ab and ACTB after knockdown of *LINC01503* by siRNA in HEK293T (left). The dot plot showing LINC01503-MP relative to ACTB in 3 independent experiments (right). The relative expression of HEK293T cells transfected with siRNA control was considered as baseline (expression= 1). siRNA control indicates the HEK293T cell treated by negative control of siRNA. ** indicates p < 0.001; one way ANOVA with Dunnett test. j. Representative western blotting by anti-LINC01503-MP Ab and ACTB in HCT116, PANC1, and U87 cells. Similar results were obtained in 2 independent experiments. k. Representative immunohistochemistry staining of healthy human kidney and colon using anti-LINC01503-MP Ab. The magnitude is 200x (left) and 100x (right). Similar results were obtained in 3 independent experiments.

Among the 72 ncORFs encoded by 66 different lncRNA genes, we focused on two overlapping ncORFs on *LINC01503*. Both ncORFs only differed in their initiation codon sites. The shorter ORF (s-ORF) is 237 nucleotides (78 amino acids; AA) in length, and the longer ORF (l-ORF) that initiates its translation 39 nucleotides upstream of the shorter isoform is 276 nucleotides (91 AA) in length (**STable1**). Both lncORFs showed high ribosome occupancy in the four tissues evaluated with Ribo-Seq (**Fig. 1d, Fig. S1, STable2**). Of note, the lncRNA, *LINC01503* itself, has been previously shown to be implicated in various cancers (Lu *et al*., 2018; Xie *et al*., 2018; Qu *et al*., 2019; M.-L. Zhang *et al*., 2020; M.-R. Wang *et al*., 2020; Shen, Sun and Xu, 2020; Ding *et al*., 2021; He *et al*., 2021; Li, Zhai and Chen, 2021; Ma *et al*., 2021; Wei *et al*., 2022; Shuai, Qian and Yuan, 2024; Weng and Huang, 2024) where it affects cell proliferation and invasion (Lu *et al*., 2018; Qu *et al*., 2019; Ding *et al*., 2021; He *et al*., 2021; Ma *et al*., 2021; Weng and Huang, 2024). However, the presence and roles of potentially encoded microproteins encoded by *LINC01503* had not been evaluated.

To summarize, Ribosome profiling revealed two ncORFs on *LINC01503*, distinguished solely by their initiation sites, and detected across multiple human tissues. This lncRNA has also been previously associated with several types of human cancers.

### *LINC01503* encodes a microprotein: LINC01503-MP

To validate the presence of the microprotein encoded by *LINC01503*, we employed parallel reaction monitoring (PRM), a targeted mass spectrometry-based proteomics assay (**see Methods**). Using this approach, we identified a unique tryptic peptide ‘FPALTQNGVALVR’ of the endogenous microprotein in human kidney tissue (**Fig. 1e, f**) and in HCT116 cells overexpressing LINC01503-MP (**Fig. S2**).

Next, we generated a rabbit polyclonal antibody (anti-LINC01503-MP Ab) targeting the C-terminal of the microprotein. This antibody successfully detected the endogenous LINC01503-MP in HEK293T cells (**Fig. 1g**). Additionally, it recognized both s-ORF- and l-ORF-derived microproteins upon overexpression in HEK293T cells (**Fig. 1g**), confirming its specificity to detect LINC01503-MP. N-terminal FLAG-tagged LINC01503-MP was also detected by anti-FLAG antibody and anti-LINC01503-MP Ab on western blotting (**Fig. S3**) and confocal microscopy analysis (**Fig. 1h**). On the other hand, C-terminal FLAG-tagged LINC01503-MP was not detected by anti-LINC01503-MP Ab (**Fig. S4**), which is likely due to the disturbance of the C-terminal epitope by the FLAG-tag. The specificity of the anti-LINC01503-MP to the C-terminal of LINC01503-MP was further validated by the siRNA-mediated downregulation of *LINC01503*, which resulted in reduced LINC01503-MP detection in HEK293T cells (**Fig. 1i**).

We further detected LINC01503-MP in other cell lines, such as HCT116 (derived from colorectal cancer) and PANC1 (derived from pancreatic cancer), but not in U87 (derived from glioblastoma) and HeLa (**Fig. 1j, Fig. S4**). In immunohistochemical staining of the kidney and colon, LINC01503-MP was mainly expressed in tubular cells of the kidney, and acinar cells of the colon mucosa (**Fig. 1k**).

Our results indicate that *LINC01503* encodes LINC01503-MP, an endogenously detectable microprotein in human tissues and specific cell lines, including cancer-derived cell lines.

### LINC01503-MP is a recently evolved, ape-specific microprotein

Evolutionary innovations can arise from alterations in translational control mechanisms and the emergence of new genes, particularly in ncORFs (Sandmann *et al*., 2023; Ruiz-Orera *et al*., 2024). Therefore, we inspected the sequence conservation of the two LINC01503-MP ncORFs by comparing the identity of the AA sequence across different primate and mammalian species. We found that LINC01503-MP exhibits poor AA conservation across mammalian species and is only conserved in apes (**Fig. 2a**). RNA-seq and Ribo-seq data supported the transcription and translation of *LINC01503* uniquely in both human and chimpanzee hearts (van Heesch *et al*., 2019; Ruiz-Orera *et al*., 2024) (**Fig. 2b**). Accumulated frameshifts and disabling substitutions across most of the mammalian species outside of the apes were found, suggesting that LINC01503-MP may have evolved unique functions or regulatory roles within primates 25 million years ago; indicating that LINC01503-MP is an evolutionarily young microprotein. Accordingly, no trace of expression of the gene and ORF was observed in the counterpart genomic region of mouse hearts (van Heesch *et al*., 2019; Ruiz-Orera *et al*., 2024) (**Fig. 2b**).

**Fig. 2.**
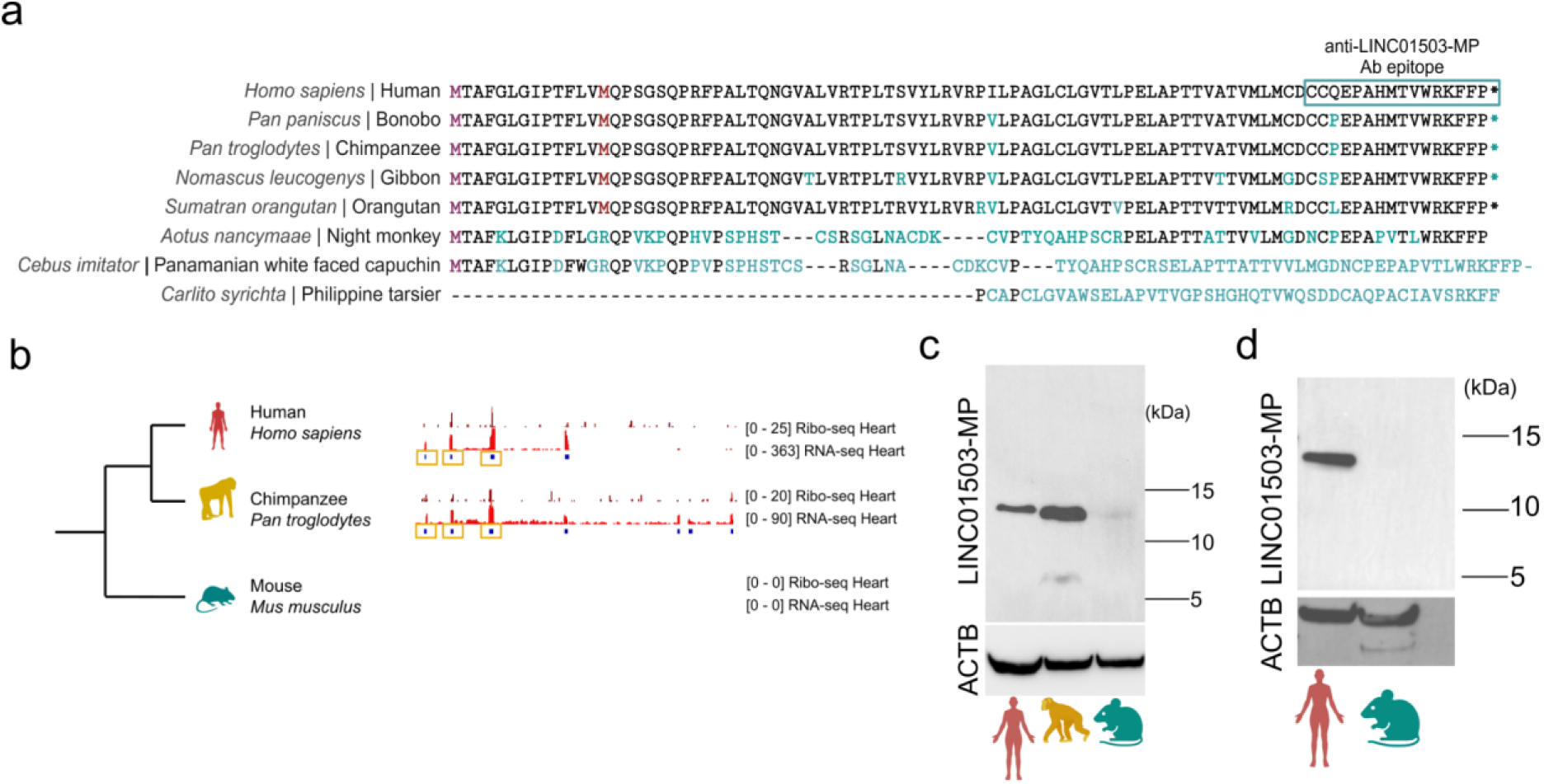
LINC01503-MP is a recently evolved, ape-specific microprotein. a. Multi-species alignment of the predicted LINC01503-MP using translated sequences from a subset of primate species retrieved from CodAlign Viewer. AA differences from humans are highlighted in blue. The putative microprotein is conserved in apes (human, bonobo, chimpanzee, gibbon, and orangutan), but displays significant frameshift substitutions in other non-ape primates (night monkey, panamanian white faced capuchin and philippine tarsier). b. RNA-seq and Ribo-seq of *LINC01503* and the encoded ncORF in human (n = 15), chimpanzee (n = 5) and mouse (n = 6) healthy hearts. Yellow boxes indicate the l-ORF encoded by *LINC01503* in humans and chimpanzees, spanning three exons. c. Representative western blotting by anti-LINC01503-MP Ab and ACTB in human, chimpanzee and mouse heart. Similar results were obtained in 2 independent experiments. d. Representative western blotting by anti-LINC01503-MP Ab and ACTB in human and mouse kidney. Similar results were obtained in 3 independent experiments.

Considering the similarity of the AA sequence of the C-terminal epitope of anti-LINC01503-MP Ab across apes (**Fig. 2a**), we speculated that the antibody may detect orthologs of LINC01503-MPs in non-human primate species. Indeed, the antibody detected a protein in chimpanzees’ hearts. Reassuringly, the microprotein was not detected in mouse hearts (**Fig. 2c**) or mouse kidneys (**Fig. 2d**), in agreement with the absence of conservation in this species.

In conclusion, these results support that LINC01503-MP represents an evolutionarily young microprotein that likely emerged *de novo* in primates, reflecting a potential unique regulatory role of this microprotein in the apes.

### LINC01503-MP localises to the mitochondria and interacts with mitochondrial proteins

Subcellular localisation may help to point to the molecular function of newly discovered microproteins and can give us clues to understand their biological roles (Sandmann *et al*., 2023). Therefore we explored LINC01503-MP’s cellular localisation using confocal microscopy. LINC01503-MP co-localised with ATPIF1, a mitochondrial protein located in the inner mitochondrial membrane (**Fig. 3a**). Importantly, the mitochondrial localisation of the LINC01503-MP was not affected by the N-terminal 3xFLAG tag (**Fig. S5**).

**Fig. 3.**
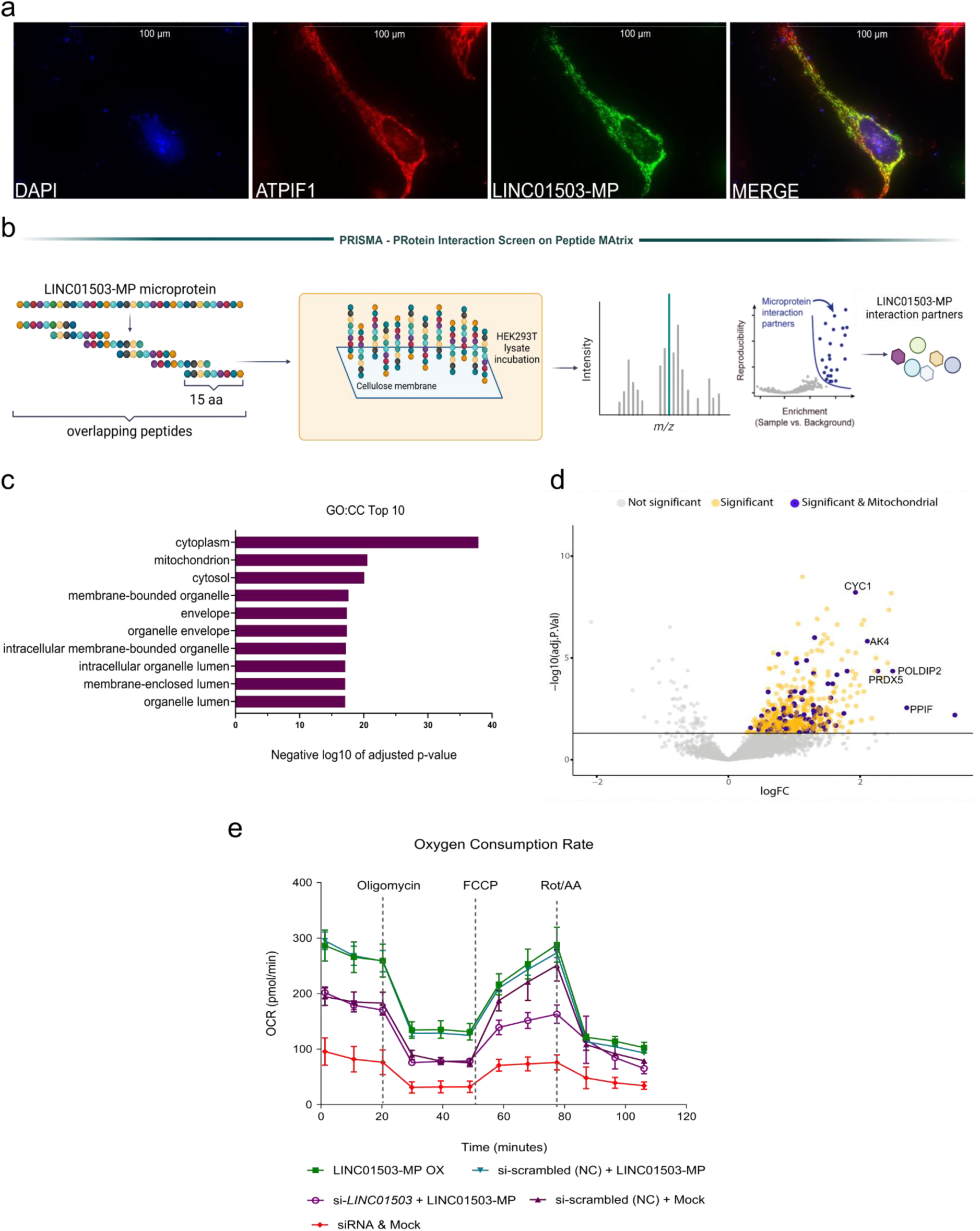
LINC01503-MP localises to the mitochondria by affecting mitochondrial function with potential interactions with mitochondrial proteins. a. Representative IF staining of overexpressed LINC01503-MP (green) co-localising with ATPIF1 (red) (a mitochondrial protein) in HeLa cells. Scale bar: 100µm. Similar results were obtained in 5 independent experiments. b. Schematic of PRISMA protocol for LINC01503-MP. c. Top 10 Gene Ontology (GO) term analysis of cellular components (CC) of all significant interacting proteins detected in PRISMA experiment. d. Volcano plot summarizing the PRISMA results of the LINC01503-MP interactome, mitochondrial interactors are highlighted in purple, including five examples involved in the electron transport chain and ATP production in the mitochondria. e. The OCR (oxygen consumption rate) profile was monitored in HCT116 cells using the Seahorse XF96 analyser. Metabolic inhibitors were injected at specific time points, as shown in the graph. Data are presented as mean ± SD (n = 5). FCCP: carbonyl cyanide-4-(trifluoromethoxy) phenylhydrazone, and Rot/AA: rotenone/antimycin A. The different groups correspond to various co-transfection conditions with controls: LINC01503-MP overexpression (LINC01503-MP-OX) is depicted in green. Co-transfection of scrambled siRNA (negative control) with LINC01503-MP-OX is shown in cyan. Co-transfection of si*LINC01503* with LINC01503-MP-OX appears in purple. Co-transfection of scrambled siRNA (negative control) with Mock (empty backbone) is represented in dark purple. Co-transfection of siLINC01503 with Mock is shown in red.

Establishing that proteins participate in specific interactions with other proteins can be crucial in gathering evidence for their potential functionality (Chen and Chen, 2021; Sandmann *et al*., 2023). Thus, to explore the potential interaction partners of LINC01503-MP, we performed protein interaction screen on peptide matrix (PRISMA) (Meyer *et al*., 2018; Dittmar *et al*., 2019; Ramberger, Sapozhnikova, *et al*., 2021; Ramberger, Suarez-Artiles, *et al*., 2021) (**Fig. 3b**). In short, we designed and synthesised 15 AA peptide fragments of LINC01503-MP with overlapping sequences. The peptides were spotted on a cellulose membrane and incubated with cell lysate for each peptide to form interactions with other proteins. Then, the binding partners of each peptide fragment were examined by mass-spectrometry. This approach not only identifies interactors but also pinpoints the specific protein regions responsible for binding. A total of 17 peptide tiles, including positive and negative controls, were synthesised and the interactome of the LINC01503-MP was established by aggregating the interactors identified in all synthesised peptide tiles. Considering the designated cut-off criteria (see **Methods**), a total of 315 significant interacting proteins were identified across all tiles, each tile exhibiting a distinct count of significant interactors (**STable 3**). We found that LINC01503-MP’s interactome was enriched in mitochondrial proteins, as well as membrane-bound organelles (**Fig. 3c**). This observation was primarily driven by a consecutive internal region (tiles 7 to 9, **Fig. 3c, STable 4**) shared by both isoforms which are predominantly engaged with mitochondrial proteins (n = 38) annotated by MitoCarta3.0 (Rath *et al*., 2021) (**Fig. 3d, STable 5**). Among the mitochondrial proteins that exhibited significant binding, CYC1, AK4, POLDIP2, PRDX5 and PPIF (**Fig. 3d**) emerged as particularly intriguing candidates among the identified interactors, given their associations with the electron transport chain and ATP production in the mitochondria. For instance, PPIF (cyclophilin D) is a protein that regulates the mitochondrial permeability transition pore (mPTP), playing a key role in apoptosis, necrosis, and mitochondrial function (Dumbali and Wenzel, 2023; Protasoni *et al*., 2024; Sautchuk *et al*., 2024).

Collectively, these results suggest that LINC01503-MP localises in mitochondria and interacts with proteins involved in mitochondrial energy metabolism.

### LINC01503-MP enhances mitochondrial respiration, rather than its associated lncRNA

To further investigate the role of LINC01503-MP in mitochondrial metabolism, a Seahorse assay (Agilent) was performed, particularly focusing on OCR (see **Methods**). After knocking down *LINC01503* by siRNA interference, we observed a significant drop in OCR levels at all phases, when compared to the controls (**Fig. 3e**) (adjusted p-value = 0.0002; one-way ANOVA corrected for multiple comparisons). Furthermore, overexpression of the LINC01503-MP showed significantly increased OCR levels compared to controls (adjusted p-value = 0.0002; one-way ANOVA corrected for multiple comparisons)in the basal respiration and proton leak. To determine whether LINC01503-MP can compensate for the phenotype induced by siRNA-mediated knockdown of *LINC01503*, we introduced an overexpression plasmid for LINC01503-MP following siRNA treatment (**Fig. 3e).** The results showed that LINC01503-MP overexpression effectively restored the OCR reduction observed upon LINC01503 knockdown (adjusted p-value < 0.0001, one-way ANOVA with multiple comparisons correction).

Collectively, our data indicate that LINC01503-MP, and not only *LINC01503* RNA transcript, contributes to regulating mitochondrial respiration, thereby supporting its involvement in mitochondrial stress responses.

### LINC01503-MP influences cell proliferation *in vitro*

Based on our findings that LINC01503-MP upregulates the levels of mitochondrial OCR, we next wanted to determine if LINC01503-MP is also involved in cell proliferation. This was motivated by previous studies showing that the *LINC01503* RNA, itself, has the potential to affect cell proliferation and invasion (Qu *et al*., 2019; He *et al*., 2021; Weng and Huang, 2024). Importantly, none of these studies evaluated the role of the ncORF and its encoded microprotein on the observed phenotypes. We therefore evaluated the impact of LINC01503-MP on cell proliferation using both siRNA knockdown and overexpression systems in HEK293T and HCT116 cell lines. This was done through a combination of cell counting to determine exact cell numbers and a CCK-8 assay to assess both cell count and metabolic activity. When we knocked down endogenous *LINC01503*, this resulted in significantly suppressed cell proliferation (p-value = 0.0014, two-way ANOVA, corrected for multiple comparisons) (**Fig 4a**). In contrast, overexpression of LINC01503-MP resulted in significantly increased cell proliferation (p-value = 0.0046, two-way ANOVA, corrected for multiple comparisons), whilst overexpression of the LINC01503-MP^△ATG^ (ncORF of LINC01503 with a mutated start codon ATG>TGA, **Fig S5**) did not have any significant effect on proliferation in number (p-value = 0.99, two-way ANOVA) (**Fig 4b**). Lastly, we performed a sequential experiment involving the overexpression of LINC01503-MP following knockdown by siRNA targeting *LINC01503*, which aimed to examine the effects of LINC01503-MP while minimizing the influence of endogenous *LINC01503* RNA and LINC01503-MP (**Fig. 4c**). The subsequent CCK-8 assay demonstrated LINC01503-MP directly affects the cell proliferation and cellular metabolism, effectively restoring proliferation inhibited by the siRNA (p-value <0.0001 for 20,000 cells and p-value = 0.0003 for 10,000 cells; two-way ANOVA) (**Fig. 4d**). Overall, these findings support that the proliferative effects previously accredited to *LINC01503* RNA in the literature may be attributed to the LINC01503-MP.

**Fig. 4.**
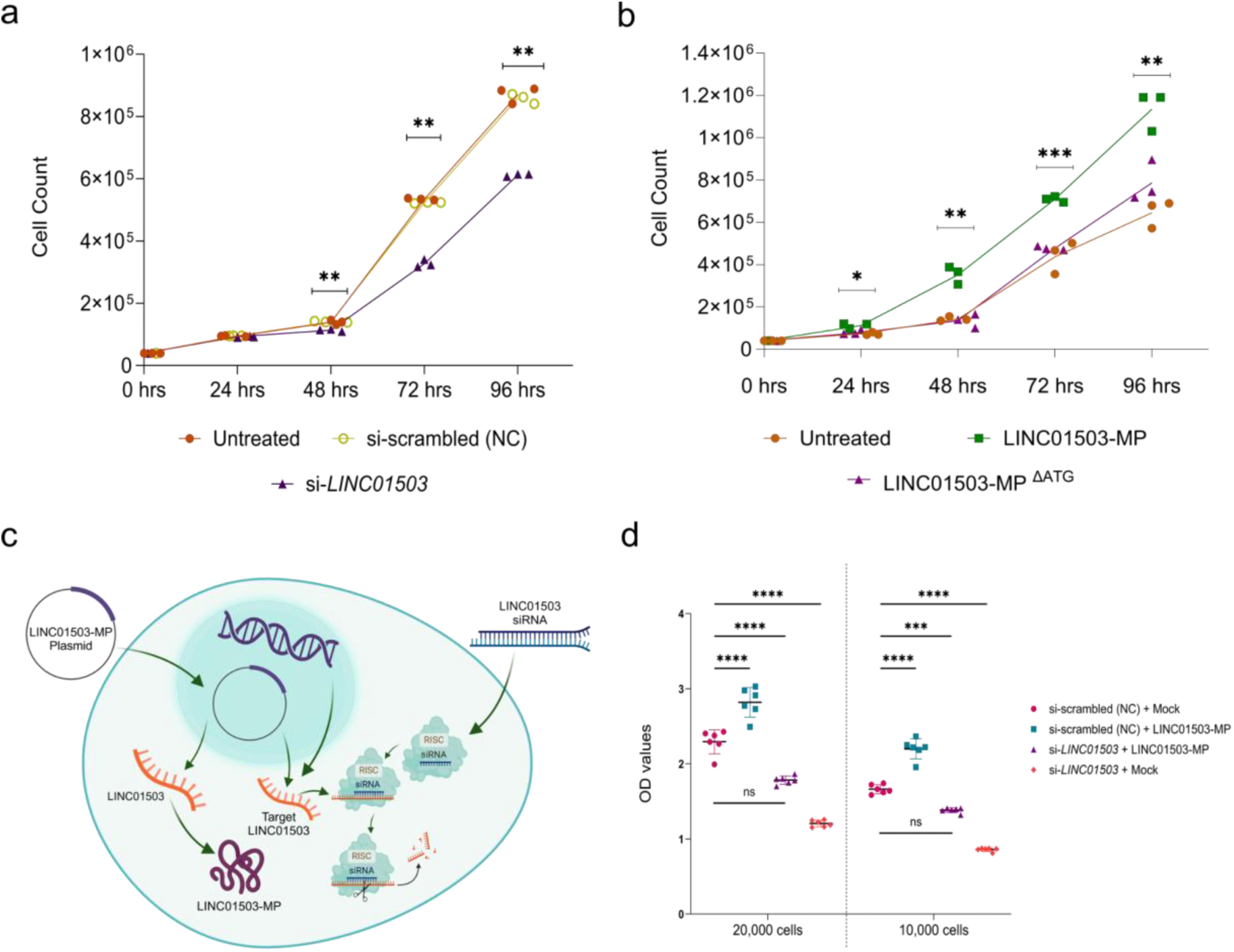
LINC01503- MP promotes cell proliferation in HEK293T cells. a. Representative cell proliferation assay in HEK293T cells after knocking down LINC01503 RNA by siRNA treatment. Similar results were obtained in 3 independent experiments. At 48 hrs, p-value - 0.0019; at 72 hrs, p-value - 0.0014; at 96 hrs, p-value - 0.0012. ** indicates p <0.01 in two-way ANOVA and Holm-Šídák tests comparing the control group and target groups were performed. b. Representative cell proliferation assay in HEK293T cells after the over-expression of LINC01503-MP or LINC01503-MP^ΔATG^. Similar results were obtained in 3 independent experiments. At 24 hrs, p-value - 0.025267; at 48 hrs, p-value - 0.006317; at 72 hrs, p-value - 0.000062; at 96 hrs, p-value - 0.020627. * indicates p < 0.05, ** indicates p <0.01, *** indicates p<0.001 in two-way ANOVA and Holm-Šídák tests comparing the control group (LINC01503-MP ^ΔATG^) and target groups were performed. c. Schematic describing the knockdown using siRNA and microprotein overexpression. d. Representative cell proliferation and cellular metabolism assay by CCK-8 assay (OD = 450 nm) in HCT116 cells. Data are presented as mean values ± SD. n = 6 technical replicates per experiment using two-way ANOVA analysis with multiple comparisons. ***P < 0.001 ****P < 0.0001 Similar results were obtained in 4 independent experiments.

### Clinical implication of LINC01503-MP in colon cancer

Previous studies have implicated *LINC01503* RNA in the proliferation of CRC (Lu *et al*., 2018; Zheng *et al*., 2024). CRC remains a significant global health challenge, with high rates of incidence and mortality (Siegel *et al*., 2023). Therefore, we investigated the expression levels of *LINC01503* in colon cancer patients compared to those without the disease. Using OncoDB (Tang, Cho and Wang, 2022; Tang *et al*., 2024), we analysed gene expression differences between colon cancer tissues and normal tissues. The pathological diagnosis of CRC samples were based on a combination of histopathological and molecular criteria, primarily using The World Health Organization (WHO) classification of tumours at the time of the study. Our analysis revealed that *LINC01503* is significantly upregulated in colon cancer samples compared to normal tissues (average transcripts per million (TPM) of 5.4 in colon carcinoma, average TPM of 2.8 in control tissue, p-value = 2.4 x 10^-9^, one-way ANOVA) (**Fig. 5a**). Additionally, the patients in the Gene Expression Profiling Interactive Analysis Database (GEPIA) (Tang *et al*., 2017) with colon carcinoma tended to show poor prognosis when *LINC01503* expression was elevated (p-value = 0.054, Mantel–Cox test) (**Fig. 5b**). Building on that, we suspected that LINC01503-MP would be also upregulated in colon carcinoma tissues. Our immunohistochemistry staining of colon tissue revealed that LINC01503-MP showed higher expression in colon carcinoma compared to control tissue (**Fig. 5c).**

**Fig. 5.**
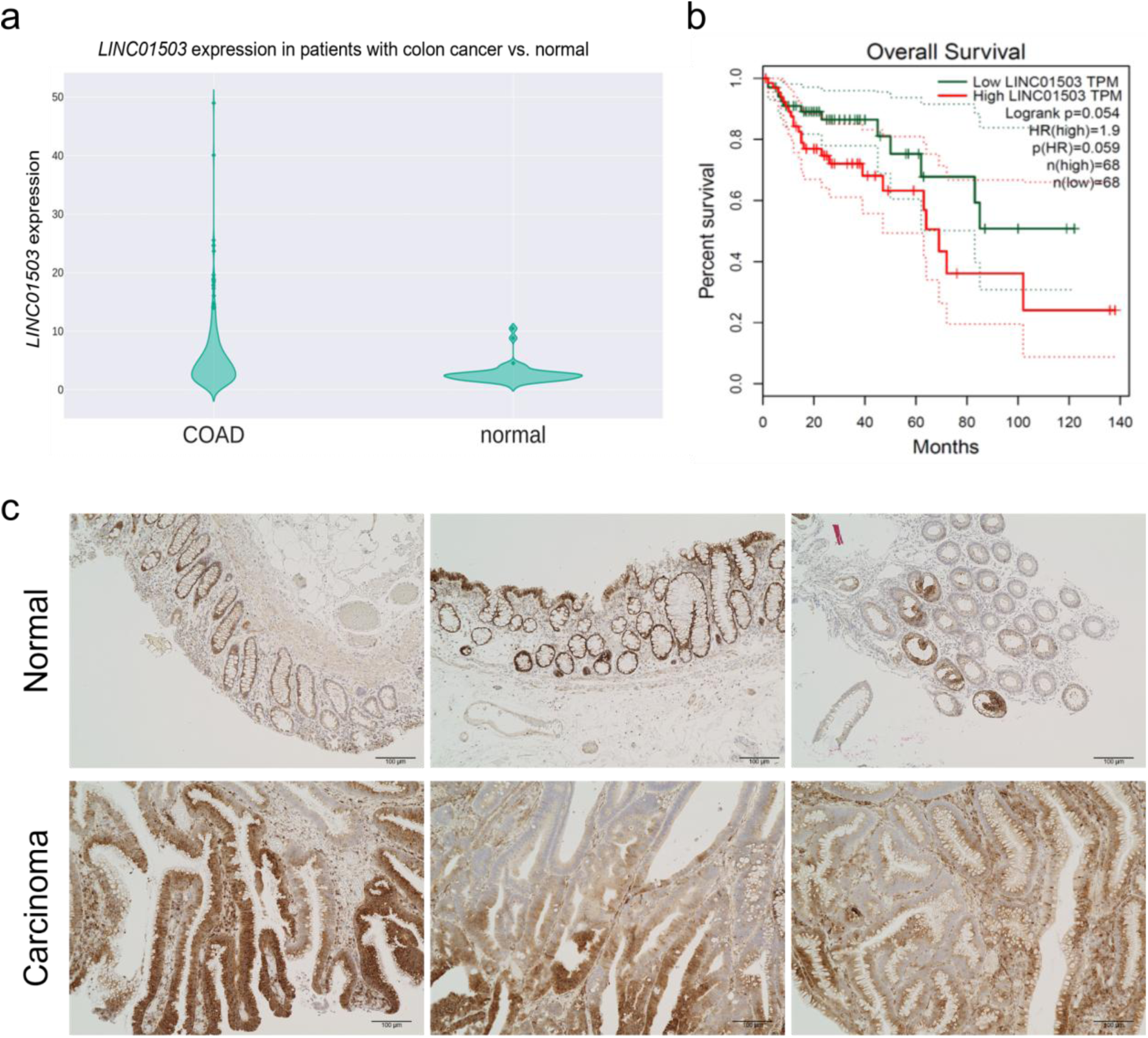
LINC01503-MP shows increased expression in colon carcinoma tissue. a. Violin plot showing the expression of *LINC01503* in patients with colon adenocarcinoma (COAD, n = 308) vs. normal (n = 41) (adapted from OncoDB) (Tang, Cho and Wang, 2022; Tang *et al*., 2024). RNA-seq data is normalised using TPM. b. Kaplan-Meier survival curve showing a tendency towards poorer prognosis in patients having higher *LINC01503* expression (adapted from GEPIA) (Tang *et al*., 2017, 2019). c. Representative immunohistochemistry staining of paraffin embedded colon carcinoma tissues (n = 3) with anti-LINC01503-MP antibody. Scale: 100 µm.

In summary, *LINC01503*, currently annotated as lncRNA, harbours a translated ncORF that produces a microprotein, which we have termed LINC01503-MP. LINC01503-MP localises in mitochondria and potentially interacts with mitochondrial proteins. LINC01503-MP can alter mitochondrial respiration and induce cell proliferation. In colon carcinoma, LINC01503-MP could relate to a poorer prognosis.

## DISCUSSION

The studies of ncORFs and their translation into microproteins has revolutionized our understanding of the transcriptome in the past decade (Ponting, Oliver and Reik, 2009; Ruiz-Orera *et al*., 2014; van Heesch *et al*., 2019). Traditionally considered non-coding genes, lncRNAs are increasingly recognized for their potential to encode biologically relevant peptides. However, the functions of most lncRNA-derived ncORFs and their translated products remain poorly understood. In this study, we identified and characterized LINC01503-MP, a novel microprotein encoded by *LINC01503*, and found that it might play a role in mitochondrial metabolism and cell proliferation based on the intracellular localisation, interactome studies (PRISMA), proliferation assays (CCK-8) and OCR analysis. In addition, we further examined its clinical implication on colon carcinoma. Similarly, literature studies have also shown that other microproteins encoded by lncRNAs are involved in cancer, for example, by promoting cell proliferation through stabilization of ATP citrate lyase, regulating ATP synthase activity, and even promoting G1/S transition by regulating cytosolic calcium levels (Pang *et al*., 2020; Ge *et al*., 2021; Yang *et al*., 2023; Zhang *et al*., 2023). Our findings, along with previous research, challenge the traditional view of lncRNAs as purely non-coding genes. Instead, they highlight the growing significance of their encoded microproteins in cellular functions and disease mechanisms.

Using ribosome profiling, proteomics and antibody-based detection such as; immunohistochemistry, immunofluorescence staining, and western blotting, we robustly identified LINC01503-MP as a microprotein translated in human tissues, including the kidney, colon, brain and heart. While the expression of the majority of lncRNAs are restricted to specific normal and/or cancer tissue types, LINC01503-MP showed consistent expression in various tissues, suggesting that LINC01503-MP, despite being involved in cancer proliferation, could also potentially exert a vital functional role across multiple organs.

Notably, LINC01503-MP does not exhibit high conservation across mammals and vertebrates like other previously described lncRNA-derived microproteins (D’Lima *et al*., 2017; Stein *et al*., 2018; Boix *et al*., 2022). We found LINC01503-MP to be an evolutionarily young microprotein that emerged during ape evolution. While most evolutionary studies only rely on sequence-based approaches (Vakirlis *et al*., 2022; Sandmann *et al*., 2023), we validated the young evolutionary origin of LINC01503-MP and its ncORF through a comprehensive approach including multiple sequence alignments, cross-species Ribo-seq data analysis, and antibody detection. It revealed that LINC01503-MP is expressed in humans and chimpanzees, but not in mice. This indicates that LINC01503-MP may contribute to ape-specific functional adaptations, as observed by previous studies indicating that ncORFs can be novel contributors to species-specific functional adaptations (Sandmann *et al*., 2023; Ruiz-Orera *et al*., 2024).

LINC01503-MP localises to mitochondria, co-localising with a mitochondrial protein, ATPIF1, located in the inner mitochondrial membrane. The interactome of LINC01503-MP revealed an enrichment in mitochondrial proteins. In addition, LINC01503-MP affects mitochondrial respiration and cell proliferation when overexpressed, suggesting LINC01503-MP may work as a mitochondrial protein cofactor to modulate cellular respiration. Interestingly, previous literature suggested that many microproteins encoded by lncRNAs are localised to the mitochondria (van Heesch *et al*., 2019; S. Zhang *et al*., 2020) and are involved in similar functions - such as regulation of mitochondrial respiratory activity; enhancing ATP synthase activity, by boosting mitochondrial ATP and oxygen use to drive colorectal cancer growth; promoting G1/S transition by regulating calcium levels and activating MAPK signaling in hepatocellular carcinoma. (Stein *et al*., 2018; Ge *et al*., 2021; Meng *et al*., 2023). Overall, these findings align with reports linking lncRNA-derived microproteins to critical metabolic processes (Cai *et al*., 2021).

Previous observations have already implicated the importance of the *LINC01503* RNA in cancer biology. *LINC01503* RNA has been shown to promote CRC progression by acting as a competing endogenous RNA for *miR-4492*/*FOXK1*, and to enhance angiogenesis in CRC by regulating VEGFA expression through *miR-342-3p* and HSP60 binding (Lu *et al*., 2018; Zheng *et al*., 2024). In this study, we demonstrated that overexpression of LINC01503-MP in HCT116 cells rescued the proliferation deficits induced by siRNA knockdown of *LINC01503*. It suggested that the microprotein might contribute to cell proliferation. In addition, we showed the high expression of LINC01503-MP in CRC compared to normal tissues using immunohistochemistry. Moreover, elevated expression of *LINC01503* RNA in CRC tissues is associated with poor patient survival. These results suggest LINC01503-MP may contribute to human cancer. However, future studies using physiologically relevant *in vivo* or *ex vivo* models would help confirm the findings obtained in human cancer cell lines.

This study has several limitations. Firstly, the endogenous expression of LINC01503-MP was not so high. It might be caused by its small size, lack of stable structure, and its instability. Due to limitations in detection efficiency, we needed to use an overexpression system to investigate the localisation of LINC01503-MP. Secondly, our study primarily relied on *in vitro* experiments, which also requires *in vivo* validation to elucidate its biological function within a physiological context. Additionally, the size of the microprotein (±9.9 kDa) detected does not match that of the band on the membrane, further post-translational modification analyses would be required to explain the size difference.

Lastly, protein interaction analysis using PRISMA required the microprotein to be synthetically generated, tiled, and spotted onto a cellulose membrane, lacking its native three-dimensional structure and physiological context, i.e. that proteins may be specific to certain cellular compartments and never have the ability to interact due to their localization in different cellular compartments *in vivo*. Therefore, the predicted interactions should be additionally validated through further assays to confirm their biological relevance.

Overall, our study establishes LINC01503-MP as an evolutionarily young microprotein translated across several human tissues, that is dysregulated in CRC and that might have a role in mitochondrial metabolism and cell proliferation. These findings suggest that the lncRNA-derived microprotein encoded by *LINC01503*, can contribute to cellular function and potentially drive cancer progression.

## DATA AVAILABILITY STATEMENT

The original contributions presented in the study are included in the article, further inquiries can be directed to the corresponding authors.

## METHODS

### Ethical statement

All experiments conducted in this study were performed under the ‘good scientific practices’ detailed by the DFG Code of Conduct.

Human kidney tissue samples were obtained at either Sapporo City General Hospital, Sapporo, Japan (ethical approval H29-057-437) or the Berlin-Brandenburg Center for Regenerative Therapies (BCRT) of the Charité Universitätsmedizin, Berlin, Germany (ethical approval EA1/134/12). The human CRC samples for the immunohistochemistry tissue staining were obtained at Sapporo Medical University, Sapporo, Japan (ethical approval 322-152).

### Collection, information, and processing of tissue samples

Snap-frozen heart tissue samples from male chimpanzees (Pan troglodytes) were supplied by the NHP tissue bank at the BPRC in Rijswijk, the Netherlands. These samples were originally collected in prior studies (Ruiz-Orera *et al*., 2024). The tissues were obtained from chimpanzees that lived at Safari Park Beekse Bergen, with necropsies conducted at the BPRC in the Netherlands. Kidney and heart tissue of the mice were obtained from C57BL/6 mice (approval number X9007/19) (van Heesch *et al*., 2019). Colon carcinoma samples were obtained from surgically or endoscopically resected specimens from five different patients.

The snap-frozen tissues were powdered using a pre-cooled mortar and pestle using constant liquid nitrogen to maintain cool temperatures. These procedures were performed on days when humidity levels were below 30%. All the technical and biological samples information were collected before obtaining and during the processing of all the samples. The tissues were lysed using RIPA Lysis and Extraction Buffer (Thermo Scientific, Waltham, USA) containing EDTA-free protease inhibitor cocktail (Roche, Basel, Switzerland) and phosphatase inhibitor (Pierce, Thermo Scientific, Waltham, USA). Then the samples were vortexed for 5 minutes and kept on ice for 1h to allow for complete lysis. The samples were then subjected to centrifugation at 14,000x*g* for 1h and pellet cell debris was removed. The lysates were then stored at -20°C until further use.

### Cell culture

The HEK293T (CRL-11268, ATCC), HCT116 (CCL-247, ATCC), HeLa (CCL-2, ATCC), PANC1 (CRL-1469, ATCC), and U87 (HTB-14, ATCC) cell lines were maintained under standard cell culture conditions in a humidified incubator at 37°C with 5% CO₂. The cells were cultured in either Dulbecco’s modified Eagle medium (DMEM) or McCoy’s medium, both supplemented with high glucose (4.5 g/L), 2 mM L-glutamine (Gibco, Thermofisher, Waltham, USA), 1 mM sodium pyruvate (Gibco, Thermofisher, Waltham, USA), or 10% foetal bovine serum (Gibco, Thermofisher, Waltham, USA). Cells were routinely passaged before reaching 80-90% confluence.

### Retrieval and Alignment of Ribosome Profiling and RNA-seq Datasets

We assembled a collection of 27 human Ribo-seq datasets from multiple studies (van Heesch *et al*., 2019; Z.-Y. Wang *et al*., 2020). These datasets were utilised to predict and quantify ncORFs across various tissues, encompassing four human organs: healthy heart (n = 15, matched RNA-seq data was also retrieved), brain (n = 6), liver (n = 3), and kidney (n = 3). We additionally analysed Ribo-seq and matched RNA-seq data for five heart samples from chimpanzees (Ruiz-Orera *et al*., 2024) and six from mice (van Heesch *et al*., 2019).

We mapped all the downloaded reads to their respective Ensembl genome (human: GRCh38/hg38, chimpanzee: Pan_tro_3.0/panTro5; mouse: GRCm38/mm38) and transcriptome (release 98) using STAR (Cardoso-Moreira *et al*., 2019) (version 2.7.3a) with the following modified parameters: *– outSAMtype BAM SortedByCoordinate*, *–outFilterMismatchNmax 2 (Ribo-seq) or 4 (RNA-seq)*, *–outFilterMultimapNmax 20*, *–alignSJDBoverhangMin 6*, *–alignSJoverhangMin 500*, *–outFilterType BySJout*, *–limitOutSJcollapsed 10000000*, *– limitIObufferSize* = *300000000* and *–outFilterIntronMotifs RemoveNoncanonical*.

### ncORF detection by ORFquant and PRICE

ncORFs were identified from the Ensembl release 98 transcriptome using our previously established pipeline (Ruiz-Orera *et al*., 2024). This approach combines ORFquant (Calviello, Hirsekorn and Ohler, 2020) (version 1.00, for AUG-initiated ORFs) and PRICE (Erhard *et al*., 2018) (version 1.0.3b, for both AUG and non-AUG ORFs), employing standard settings and including only uniquely mapped reads. Significant ORFs (adjusted p-value < 0.05) identified by either software in each sample were merged to create a comprehensive list of translated ncORFs in lncRNAs. ORFs with ≥ 90% sequence identity were grouped, prioritising AUG-initiated ORFs, and the longest ORF in each group was selected as the representative. For each Ribo-seq sample, we used RiboseQC to extract P-site counts, which were used to quantify in-frame P-site counts for each ncORF. The Integrative Genomics Viewer (IGV) (Robinson *et al*., 2011) was used to visualize RNA-seq, Ribo-seq, and P-site coverage for the genomic loci of human and chimpanzee *LINC01503*, as well as the orthologous region in mice. For visualization, all samples per organ were pooled to generate combined coverage tracks.

### Mass spectrometry analysis and tryptic peptide mapping

#### Protein Extraction and Digestion

Cell lysates were lysed in 2% SDS-DTT buffer (10 mM dithiothreitol [DTT], 100 mM Tris-HCl, pH 8.0). The lysate was diluted with SDS-DTT buffer to a final concentration of 1% SDS, 5 mM DTT, and 50 mM Tris-HCl, pH 8.0. Samples were vortexed, briefly centrifuged, and incubated at 90°C for 10 min to denature proteins. For protein alkylation, iodoacetamide (IAA) was added to a final concentration of 10 mM, and the samples were incubated for 30 min at 25°C in the dark. The reaction was quenched by adding DTT to a final concentration of 20 mM. The pH was verified to be neutral before proceeding. Proteins were precipitated using a bead-based approach. For SP3 cleanup and digest (Hughes *et al*., 2014), Sera-Mag Speed Beads, CAT# 09-981-121, and CAT# 09-981-123 (Thermo Fisher Scientific) 1:1 bead mix was added at a final concentration of 100 µg/µL (1:10), followed by the addition of acetonitrile (ACN) to achieve a final concentration of >70%. Samples were incubated at room temperature for 18 min to allow protein binding. Beads were then pelleted using a magnetic rack, and the supernatant was removed and discarded. Beads were washed three times with 200 µL of 70% ethanol. Bound proteins were resuspended in 100 µL of digestion buffer containing 100 mM HEPES, pH 8.0, Lys-C (0.5 mAU/µL), and trypsin (0.5 µg/µL) at a 1:25 enzyme-to-protein ratio. Proteins were digested overnight at 37°C with agitation (1150 rpm) in a thermomixer. Following digestion, the beads were separated on a magnetic rack, and the supernatant containing peptides was transferred to fresh tubes. Beads were rinsed with 100 µL of 100 mM HEPES, pH 8.0, and the rinse was combined with the previous supernatant. Peptides were acidified by adding 20 µL of 10% formic acid (FA), ensuring the pH was below 3. The peptide solution was desalted using the AssayMAP Bravo Protein Sample Prep Platform (Agilent Technologies) according to the manufacturer’s protocol.

#### Parallel reaction monitoring (PRM) analysis

Stable isotope-labelled synthetic peptides were synthesised in the SpikeTides TQL format (JPT Peptide Technologies) with the following sequences: {Ac-TAFGLGIPTFLVMQPSGSQP-R*-Qtag} and {H-FPALTQNGVALV-R*-Qtag} where the asterisk indicates a heavy arginine 10 and the Ac an acetylated peptide N-terminal. Each peptide was solubilised separately in 100 µL of solubilisation buffer (8 M urea, 50 mM Tris-HCl, pH 8.0) to a final concentration of 10 pmol/µL. Peptides were sonicated for 5 min in a water bath to ensure complete dissolution. For digestion to remove the Qtag, 100 µL of each peptide solution (containing 1 nmol of peptide) was treated sequentially with 12 mM DTT for 30 min at 32°C, followed by alkylation with 40 mM IAA for 30 min at 25°C in the dark. Proteins were then digested overnight with 0.5 µg of sequencing-grade trypsin (Promega) at 37°C. The reaction was acidified by adding 1% FA, and peptides were desalted using StageTips (Rappsilber, Ishihama and Mann, 2003). Desalted peptides were dried and resuspended in MS sample buffer (3% acetonitrile, 0.1% FA). The peptides were then pooled and diluted to a final concentration of 100 fmol/µL in the MS sample buffer. For LC-MS analysis, 2 µL of peptide solution, corresponding to 100 fmol of synthetic peptide and 500 ng of digested sample, was injected. Samples were analysed using an Orbitrap Astral mass spectrometer (Thermo Fisher Scientific) in tMS² mode, coupled to a Vanquish Neo nano-LC system (Thermo Fisher Scientific) for peptide separation using essentially the same LC-method as described for the DIA experiment. Mass spectrometry was performed in positive ion mode using a Nanospray Flex Ion Source (Thermo Fisher Scientific) with a source voltage of 2.2 kV and an ion transfer tube temperature of 280°C. MS1 scans were acquired at a resolution of 240,000, with a scan range of 380–1100 m/z, RF lens setting of 40%, an AGC target of 500%, and a maximum injection time of 3 ms. tMS² scans were acquired with an isolation width of 1.3 m/z. Precursor ions were selected based on their m/z values and charge states, with optimised higher-energy collisional dissociation (HCD) normalised collision energy settings applied individually. The light and heavy-labelled versions of T(+42.010565)-AFGLGIPTFLVM-(+15.994915)-QPSGSQPR were monitored at m/z 754.7244 (z = 3, 40% HCD) and m/z 758.0605 (z = 3, 40% HCD), respectively. Similarly, FPALTQNGVALVR was monitored in two charge states: m/z 462.6015 (z = 3, 30% HCD) and m/z 693.3986 (z = 2, 30% HCD) for the light form, and m/z 465.9376 (z = 3, 30% HCD) and m/z 698.4028 (z = 2, 30% HCD) for the heavy form. MS2 spectra were recorded within a scan range of 150–2000 m/z.

Raw data were processed using Skyline (v24.1.0.199; MacLean, 2010) for targeted peptide quantification. Annotated MS2 spectra were generated with MaxQuant (v1.6.3.4; Cox et al., 2008) following peptide identification via a database search against the UniProt human database (release 2022-03), including isoforms, the putative LINC01503-MP sequences, and common contaminants. Arginine-10 was set as a variable modification alongside oxidation (M) and N-terminal acetylation. Spectra were visualized and exported using MaxQuant Viewer (Rappsilber, Mann and Ishihama, 2007; Cox and Mann, 2008; MacLean *et al*., 2010; Hughes *et al*., 2014).

#### Peptide Mapping

Using ProteoMapper (Mendoza *et al*., 2018) we mapped the tryptic peptides to the human peptide atlas. The tryptic peptide ‘FPALTQNGVALVR’ was unique to the LINC01503-MP sequence and ha a PSS score (Predicted suitability score, derived from combining publicly available algorithms [Peptide Sieve, STEPP, ESPP, APEX, Detectability Predictor]) of 0.93.

### Protein sequence conservation of ncORFs on *LINC01503*

We used CodAlignView (https://data.broadinstitute.org/compbio1/cav.php) to check the sequences of the human ncORF on *LINC01503* against 60 primates and manually identified key substitutions that truncated the ncORF in other non-human species. With exon intervals with alignment set ‘hg38_470mammals_primates’ on chromosome 9 and as follows; ‘chr9:129336977-129336988+chr9:129338792-129338917+chr9:129341736-129341873’.

### Plasmid generation

The s-ORF and l-ORF (**STable 6**) were synthesised by Integrated DNA Technologies (IDT, Coralville, USA) with or without N-terminal FLAG-tag. The synthesized nucleotides were inserted to p3xFLAG-CMV-14 (Sigma-Aldrich) using restriction enzymes BamHI and EcoRI (New England Biolabs, Massachusetts, United States).

To disrupt the predicted ncORF, the Q5^®^ Site-Directed Mutagenesis Kit (New England Biolabs, Massachusetts, USA) was used to mutate the start codon, AUG to TGA. Mutagenic primers (**STable 7**) were designed using the NEBaseChanger tool (https://nebasechanger.neb.com/), and they were synthesised and HPLC purified by BioTeZ (Berlin, Germany). Mutagenic PCR reactions, DpnI digests, and bacterial transformation were performed according to the manufacturer’s instructions. The plasmid DNA was extracted from transformed bacterial cells using QIAGEN miniprep & maxiprep kits (QIAGEN, Hilden, Germany). The inserted and insertion sites sequences were verified by Sanger sequencing (Azenta Life Sciences, Massachusetts, USA). The plasmid construct information is in **STable 8**.

### Custom antibody generation for LINC01503-MP

The estimated C-terminal peptide sequence (15 AA) of LINC01503-MP was synthesised and immunised four times every 2 weeks to *Oryctolagus cuniculus*, and the whole serum was collected. The serum was performed affinity purification by the epitope antigen. The antibody, named anti-LINC01503-MP Ab, was manufactured by Sigma Aldrich, Tokyo, Japan. The purified antibody was stored -80°C until further use.

### Transfection of plasmids

Cells were transfected with 5 µg of plasmid per well (growth area 9.07 cm²) using LipoJet transfection reagent and buffer according to the manufacturer’s protocol (LipoJet™ In Vitro Transfection Kit (Ver. II) SignaGen Laboratories, Rockville MD, USA). Cells were then incubated at 37°C at 5%CO2 for two days. The transfected cells were further used for western blotting, cell proliferation assay or immunoprecipitation.

### SDS-PAGE and western blotting

Sample lysates (11 µL) were mixed with 4 µL NuPAGE™ LDS Sample Buffer (4X) (Thermofisher Scientific, Waltham, USA) and 1.6 µL NuPAGE™ Sample Reducing Agent (10X; Thermofisher Scientific, Waltham, USA) and incubated at 70°C for 10 minutes. Proteins were separated by SDS-PAGE on NuPAGE™ 12% Bis-Tris gels (Thermofisher Scientific, Waltham, USA) in a 1x MES buffer (Thermofisher Scientific, Waltham, USA) at 180 V for 45 minutes at a constant voltage setting. Protein size was determined using Precision Plus Protein™ Dual Xtra Prestained Standards (Bio-Rad California, USA).

For western blotting, PVDF membranes (Immobilon-PSQ) 0.2 µM (Merck Millipore, Massachusetts, USA) were activated in ethanol (VWR, Pennsylvania, USA), washed with transfer buffer (1x NuPAGE™ transfer buffer; Thermofisher Scientific, Waltham, USA) with 20% ethanol), and equilibrated in transfer buffer. A blotting sandwich consisting of filter paper, membrane, gel, filter paper, and sponge was subjected to tank blotting at 90 mA, 4°C for 1.5 hours in constant current setting.

Following the transfer, membranes were washed with TBS-T (1x TBS, 0.1% Tween) and blocked for 1 hour with 10% BSA in TBS-T at room temperature. The membranes were incubated overnight at 4°C with primary antibodies (**STable 9**) in TBS-T containing 5% BSA. After washing three times with TBS-T, the membranes were incubated for 1 hour with HRP-conjugated secondary antibody (**STable 9**) in TBS-T containing 5% skim milk. The membranes were washed twice with TBS-T and HRP-mediated chemiluminescence was detected using a Bio-Rad ChemiDoc imager following the application of a chemiluminescence (ECL) reagent (Amersham Biosciences, Amersham, UK). The membrane was cut at the 25 kDa line and then stained with the housekeeping protein or, after stripping the membrane with western blot stripping buffer (Santa Cruz, Texas, United States) following manufacturer’s instructions, then the membranes were reprobed with monoclonal anti-ACTB Ab (**STable 9**).

### Knockdown by siRNA

siRNAs were designed based on two previous publications (Lu *et al*., 2018; Ma *et al*., 2021). They were synthesised from IDT at 10 nmoles total concentration. The Scrambled Negative Control DsiRNA (5 nmol, IDT) were used as negative controls. The control does not recognize any sequences in human, mouse, or rat transcriptomes. All siRNAs were suspended in nuclease-free water to a stock concentration of 100 µM and then later to a working concentration of 10 µM resuspended in distilled nuclease-free water, and stored at -20°C until further use. 400,000 HEK293T cells/well were cultured in standard tissue culture plates -6 well (Sarstedt, Germany) for 24h and transfected with a pool of 10 µM siRNA or scramble siRNA in each well using Lipofectamine™ RNAiMAX (Thermofisher, Waltham, USA) reagent following manufacturer’s instructions and were incubated for 48h. Post transfection, the cells were washed twice with ice-cold phosphate-buffered saline (PBS), and scraped and lysed in 200 µL lysis buffer. After centrifugation at 14,000x*g* for 40 min at 4°C, the supernatant of the lysate was used for further analysis to remove the cell debris.

Images of the western blot membrane were captured using ChemiDoc imager (Bio-Rad) ensuring consistent exposure settings and resolution across all samples. The acquired images were converted to 8-bit grayscale images using Fiji (ImageJ). For each lane, the target band and a corresponding housekeeping protein band were identified in the region of interests by rectangle selection and quantified the integrated density values of each band. The intensity values of the target protein bands were normalized to their corresponding housekeeping protein bands.

### Immunohistochemistry (IHC)

The obtained tissue was fixed by 10% formaldehyde and embedded in paraffin. The paraffin block was sectioned to 5 um and attached onto glass slides. The tissue slides were further treated by an automated IHC system (BOND-MAX, Leica, Japan). The slides were deparaffinized, and rehydrated. Antigen retrieval was performed by BOND-Epitope Retrieval Solution 2 (EDTA based pH 9.0, Leica, Japan) with heating (100°C, 20 min). To block the endogenous peroxidase activity, the slides were treated by 3% hydrogen peroxide. Blocking was performed by block ace solution (WakenBtech Co., LTD. Japan) for 20 min. Primary antibody reaction (**<u>STable 9</u>**) was performed for 15 min and secondary antibody reaction was 8 min. 3, 3-diaminobenzidine (DAB) staining was performed by Bond Polymer Refine Detection (Leica, Japan) for 10 min. Counter staining was performed by Haematoxylin for 4 min. The images were obtained by Olympus BX40 microscope (Olympus, Japan). (number of patients = 5) (n=3 representative data shown in Fig 5)

### Immunofluorescence (IF) Imaging

HeLa cell line, which have very low endogenous expression of LINC01503-MP, were used with over-expressing LINC01503-MP or N-terminal 3xFLAG fused LINC01503-MP. The transfected cells were plated onto µ-Slide 8 Well high chamber slides (ibiTreat, Ibidi) 24 hours before the experiment and cultured in the cell incubators. The cells were fixed with 4% paraformaldehyde (Sigma-Aldrich), washed with PBS, and incubated with a blocking and permeabilizing solution containing 10% normal goat serum (Gibco) and 0.1% Triton X-100 (Sigma-Aldrich) in PBS with 0.05% Tween 20 (Cell Signalling) and then washed twice again after the incubation. LINC01503-MP was stained overnight at 4°C using the anti-LINC01503-MP Ab (**STable 9**) or anti-FLAG Ab (**STable 9**) and co-stained with ATPIF1 (**STable 9**). The chamber-slides were washed twice and incubated with fluorescently-labelled secondary antibodies (**STable 9**) for 1 hour in the dark at room temperature. The cells were washed again with PBS and stained with 4-6-diamidino-2-phenylindole (NucBlue Fixed Cell ReadyProbes Reagent, R37606, Thermo Fisher) for 10 minutes at room temperature. And then Fluoromount G (Invitrogen) was used to mount the cells in the chamber slides. Images were visualized using a LEICA SP8 confocal microscope using a 63x/1.30 Glycerol HC PL Apo CS2. The images were acquired with an inverted confocal laser scanning microscope Leica TCS SP8, with the LAS X (v. 3.5.5) acquisition software. Imaging was performed with a scan speed of 400 Hz (pixel dwell of 0.2 µs), unidirectional scan direction X, the pinhole set to 1AU, a pixel size 60x60 nm, a line average of 4, and excitation beam splitter TD 488/561/633. Image analysis was performed using FIJI (V2.9.0) (Schindelin *et al*., 2012).

### PRISMA

#### Design of membrane

The PRISMA experiment described in this study was adapted from a previously established assay documented in the literature (Dittmar *et al*., 2019; Ramberger, Suarez-Artiles, *et al*., 2021). LINC01503-MP was divided into 13 mini-overlapping peptide sequences, referred to as “tiles.” Each tile consisted of 15 AA peptides in length, with a total of 8 AA overlap between adjacent tiles. These peptides per tile were individually synthesised using SPOT synthesis technology on a cellulose membrane provided by JPT Inc., Berlin, Germany (**STable 10**). Each spot contained approximately 5 nmol of covalently bound mini peptide, attached to the cellulose-ß-847 alanine-membrane. The synthesis of control peptides, SOS1 and GLUT1, was also performed on separate spots on the same membrane, following previously reported procedures (Meyer *et al*., 2018) (Schulze and Mann, 2004).

#### Protein lysate preparation and incubation with membrane

HEK293T cells were cultured and grown in 15 cm dishes (Sarstedt). Throughout the experiment, all procedures were conducted on ice (∼4°C) using exclusively ice-cold buffers. The cells were washed with PBS with calcium and magnesium (PBS^+/+^) and subsequently scraped using a cell scraper. The cell lysate was collected and centrifuged for 5 minutes at 1000 xg. Following an additional wash with PBS^+/+^, the cell pellets were resuspended in lysis buffer (50 mM HEPES pH 7.6 at 4°C, 150 mM NaCl, 1 mM EGTA, 1 mM MgCl2, 10% Glycerol, 0.5% Nonidet P-40, 0.05% SDS, 0.25% sodium deoxycholate, and cOmplete™ EDTA-free protease inhibitor (Roche)). The re-suspended lysate (0.7 mL per 14 cm dish) was then incubated for 30 minutes on ice. Following the incubation, 5 μL (1250 units) of benzonase (Merck) was added, and the mixture was further incubated for 15 minutes before being centrifuged for 15 minutes at 20,000x*g*. The resulting supernatant was transferred to a fresh falcon tube, and the protein concentration was determined using the Pierce™ BCA Protein Assay Kit (Thermofisher Scientific), following the manufacturer’s protocol. The protein concentration was adjusted to 5 mg/mL using the lysis buffer. This protein extract was directly used for the subsequent incubation with the cellulose membrane. The cellulose membrane carrying the synthesised mini peptides was first hydrated for 15 minutes in a wash buffer at room temperature (RT) on a shaker. Subsequently, it was incubated with a 1 mg/mL tRNA blocking buffer (Invitrogen) diluted in wash buffer for 10 minutes, followed by two washes with wash buffer for 5 minutes each. Next, the membrane was incubated with the cell lysate (5 mg/mL) for 2 hours at 4°C while gently shaking. Following the incubation, the membrane was washed three times with a wash buffer for 5 minutes at 4°C and then dried for 15 minutes at RT. After the incubation and drying steps, the peptide spots were carefully punched out using a 2 mm puncher and transferred directly into 20 μL of urea sample buffer (6 M urea, 2M thiourea, 10 mM HEPES-KOH, pH 8.0). To prepare the samples for further analysis, they were first reduced by incubating in a 12 mM DTT for 30 minutes at RT. Subsequently, alkylation was performed by treating the samples with 40 mM chloroacetamide for 45 minutes at RT in the dark. For protein digestion, the samples were diluted with 100 μL of a 50 mM ammonium bicarbonate buffer (pH 8.5) containing trypsin (Promega; 5 μg/mL) and LysC (Wako; 5 mAU/mL). The diluted samples were then incubated overnight at RT to allow for proteolytic digestion. The proteolytic digestion was stopped by adding 4 μL of 25% trifluoroacetic acid, and the peptides were subsequently extracted and desalted using the Stage-Tip protocol with two disks of C18 material (Rappsilber, Ishihama and Mann, 2003).

#### LC-MS/MS

Preparation for MS (i.e., incubation of the membrane with HEK239T protein lysate, punching out the peptide spots, digesting the peptide spots with trypsin and LysC, preparation of the Stage Tips), LC-MS/MS (i.e., elution of the peptides, separation of the peptides on an HPLC system, ionisation, and analysis of the peptides on an Orbitrap Fusion instrument) as well as raw data analysis and filtering steps for identification of protein-protein interactions were performed as described in the above PRISMA screen for microproteins.

#### Data analysis and quality control

The raw data obtained from the experiment were subjected to peak detection and analysis using MaxQuant software (version 1.6.3.4). The MS data was searched against the human UniProt database (2022-03) and an in-house database containing the microprotein sequence information. LFQ intensities were log2-transformed and kept as is or normalized using median-MAD normalization, i.e. {xi - median(x))/mad(x)}. This is similar to a z-score which is {xi - mean(x))/sd(x)} in a column- (=experiment) wise manner.

Downstream analysis was performed in R using the proteinGroups.txt output from MaxQuant. Contaminants were filtered out from the protein list as well as ribosomal proteins which were identified as a common contaminant in the pulldowns. LFQ intensities were log2-transformed and additionally normalised by median-MAD scaling. Only protein groups with 3 valid values for each experimental group to be compared were selected. For each pairwise comparison, imputation of remaining missing was performed using a randomised Gaussian distribution with downshift (shift=1.8; width=0.3). Two-sample moderated t-tests were performed for the group-wise comparisons using the limma package (4). Each experiment was compared to a blank sample (empty pulldown) as well as the controls, i.e., all the other pulldowns combined. For the LINC01503-MP comparisons, LINC01503-MP_CQEPAHMTVWRKFFP was excluded from the controls. This is because the pulldowns had less valid values but higher intensity distribution. Furthermore, GLUT1 and SOS1 peptide pulldowns served as positive controls for the assay’s performance (3). Proteins with an adjusted p-value ≤ 1e-4 and a logFC ≥ 1.5 were considered as significant candidates.

### Gene ontology analysis

The gene ontology (GO) enrichment analysis for the LINC01503-MP interactome identified with PRISMA was performed with gProfiler2 v0.2.0 (Kolberg *et al*., 2023) with default parameters.

### Cell proliferation assay

#### Cell counting

HEK293T cells were grown and transfected as described above. 24 hours post transfection, the cells were harvested and mixed with equal parts of 0.4% Trypan Blue (Thermofisher, Waltham, USA) solution and pipetted into a Countess chamber slide and then inserted into Invitrogen Countess 3 Automated Cell Counter (Thermofisher, Waltham, USA). The function “Rapid Capture” was used to count the cells.

#### CCK-8 assay

On Day 0, HCT116 cells were resuspended at 400,000 cells/mL in Falcon or Eppendorf tubes. On Day 1, reverse transfection was carried out using siRNAs targeting *LINC01503* or scrambled siRNA with siQuest reagent in Opti-MEM, and cells were plated in 6-well plates at 500,000 cells/well. Simultaneously, cells were transfected with LINC01503-MP overexpression or mock plasmids. On Day 2, cells were harvested, counted, and diluted to 18,000 cells/100 µL or 9,000 cells/100 µL before being plated in 48-well plates (300 µL/well). A calibration curve was generated using known cell numbers, and plates were incubated overnight at 37°C, 5% CO₂. CCK-8 reagent was thawed and added to each well (10 µL/well), followed by a 3-hour incubation at 37°C. Absorbance was measured at 450 nm, with background absorbance recorded at 650 nm. A calibration curve was generated, and plates were preserved with 10 µL of 1% SDS for future measurements.

Cell viability was calculated as follows:

- Cell Viability (%) = (Absorbance of treated cells−Background absorbance) (Absorbance of control cells−Background absorbance) × 100

The experiments were conducted in technical replicates of 6 and biological replicates of 3, with results normalized to control conditions.

### Seahorse assay

The seahorse assay was performed with HCT116 cells using the Agilent XFe96 Analyser instrument (software used: WAVE 2.4.3). The protocol was followed as per manufacturer’s instructions using the Seahorse XF Cell Mito Stress Test Kit, by Agilent Technologies (Cat#103015-100). Treated HCT116 cells were subjected to OCR measurements at 37°C in an XF96 extracellular flux analyser (Agilent Technologies). The Seahorse XF96 Cell Culture Microplate (Cat#101085-004) was used to seed and culture the HCT116 cells (4×10^4^). The treated cells well seeded into the Seahorse XF96 cell culture microplate, and plated in DMEM media pH 7.4 that was supplemented with 25 mM glucose and 1 mM sodium pyruvate and sequentially exposed to oligomycin (2.5 μM), FCCP (1.0 μM), and Rot/AA (0.5 μM) as previously reported in (Luongo *et al*., 2015) and from the manufacturer’s recommended doses.

### Database analysis

The patient survival analysis plot was generated using GEPIA (Tang *et al*., 2017, 2019) using the quartile group cutoff. The cutoff determines how the expression levels of the LINC01503 gene are divided into groups for Kaplan-Meier survival analysis. The dataset used is split into high-expression (top 25%), low-expression (bottom 25%). Since it matters if *LINC01503* expression levels are high or low, we used the quartile cutoff; this approach takes into account the threshold-dependent effect (e.g., very high or very low expression). The GEPIA web server uses log-rank tests, also known as the Mantel–Cox test, for the hypothesis evaluation. The database GEPIA uses is the TCGA and GTEx gene expression data.

### Quantification and statistical analyses

The siRNA and overexpression data were expressed as the mean ± standard deviation (SD) of the normalized band intensities from at least three independent experiments. Statistical significance between experimental groups was determined using either one-way ANOVA or two-way ANOVA, with a significance threshold set at p < 0.05 using Graphpad Prism software v10.2.2. The data were visualized using dot plots using Graphpad Prism software.

Statistical analyses and normalisation for the PRISMA experiment was done using a two-sample moderated t-test, and significance based Benjamini-Hochberg (BH) post hoc test. We also performed multiple-comparison correction using the BH-procedure resulting in the adjusted p-values.

*For each comparison*:

Only rows (=proteins) containing 3 valid values for the experimental group were selected. The remaining selected control columns were imputed using a randomized Gaussian distribution with downshift {rnorm(length(x[is.na(x)]), mean(x)-shift*sd, width*sd)} with;

- sd= standard deviation by column
- shift=1.8; width=0.3

*A two-sample moderated t-test was applied between groups*:

1. Experiment vs. BLANK
2. Experiment vs. controls
3. Using the limma package with the parameters {robust=TRUE, trend=TRUE}

GraphPad v10.2.2 was used to generate plots and to perform two-tailed t tests or one-way ANOVAs with either Dunnett’s or Bonferroni post hoc to obtain the reported p values. All results were considered to be significant if p values after correction were ≤ 0.05 (false discovery rate [FDR] 5%). Sample numbers are indicated in the figure panels or legends. Reproducibility was ensured by having biological replicates per condition/treatment.

## ACKNOWLEDGEMENTS

The authors thank the Advanced Light Microscopy Technology Platform at MaxDelbrück-Center for Molecular Medicine in the Helmholtz Association (MDC), Berlin, Germany (https://www.mdc-berlin.de/advanced-light-microscopy) and especially Dr. Sandra Cristina Carneiro Raimundo for her technical support and assistance with confocal microscopy imaging in this work. M.K. was supported by JSPS KAKENHI Grant Number 20K22906 and 21K15979. M.K. received funding provided by the Alexander von Humboldt Foundation.

N.H. was supported by a grant from the Leducq Foundation; an ERC Advanced Grant under the European Union Horizon 2020 Research and Innovation Program (AdG788970); a British Heart Foundation and a Deutsches Zentrum für Herz-Kreislauf-Forschung grant (BHF/DZHK: SP/19/1/34461); the German Research Foundation (DFG) (CRC/SFB-1470 – B03); and, in part, by a grant from the Chan Zuckerberg Foundation.

S.v.H. was supported by a DZHK (German Centre for Cardiovascular Research) Excellence Program PostDoc Start-up Grant (81X3100106), and Oncode Institute, which is partly financed by the Dutch Cancer Society (KWF). This publication is part of the project “Evolutionarily young microproteins in childhood brain cancer” (with project number VI.Vidi.223.022 of the research programme NWO talent programme Vidi, which is (partly) financed by the Dutch Research Council (NWO), awarded to S.v.H.

This work was supported by the Deutsche Forschungsgemeinschaft (DFG—German Research Foundation) under grant agreement SFB 1470 “HFpEF” (Project B05) and SFB 1588 “Neuroblastoma Evolution” (Project A06) to P Mertins.

## AUTHOR CONTRIBUTIONS

Conceptualization and methodology: N.D; M.K; S.v.H; N.H

Validation: N.D; JR-O; O.P; M.K

Formal analysis: N.D; JR-O; F.W; C.S O.P; M.K

N.D performed most experiments with some assistance from M.K; M.S; O.P; S.B; N.L; H.N

Data curation: N.D; JR-O; O.P; M.K

Resources: N.H; M.K; JR-O; O.P; M.S; JF-S; T.T; I.K; S.O; S.Y; H.K; A.K. H.N; S.v.H; P.M

Writing – original draft: N.D; JR-O; O.P; M.K

N.D; JR-O; M.K critically revised the manuscript; with input from all authors M.S; N.L; I.K; O.P; S.B; H.K; T.T; S.O; P.M; N.H; S.v.H; JF-S; F.W; S.Y; A.K; H.N; C.S; T.K

Supervision: JR-O; M.K; N.H; P.M

Funding acquisition: N.H; P.M; M.K

## DECLARATION OF INTERESTS

The authors declare no competing interests.

## Notes

### Competing Interest Statement

The authors have declared no competing interest.

