## Supplementary Information for "LINC01503-MP is a Mitochondrial Microprotein That Promotes Cell Proliferation and Oxidative Metabolism"

#### Supplemental Information

### LINC01503-MP is a Mitochondrial Microprotein That Promotes Cell Proliferation and Oxidative Metabolism.

Nikita Dewani[1,2], Jorge Ruiz-Orera[1], Oliver Popp[3], Ning Liang[1], Masanari Sugarawa[4], Jana F. Schulz[1], Franziska Witte[1], Clara-Louisa Sandmann[1], Takahiro Tsuji[5], Susanne Blachut[1], Takaharu Katagiri[1], Ivanela Kondova[6], Sae Owada[7], Shinji Yoshii[7], Hiroshi Kataoka[8], Andreas Kurtz[9], Hiroshi Nakase[7], Sebastiaan van Heesch[10,11], Philipp Mertins[3,13], Norbert Hübner[1,2,12,13,14,#], Masatoshi Kanda[1,4,14,#]

[1] Cardiovascular and Metabolic Sciences, Max Delbrück Center for Molecular Medicine in the Helmholtz Association (MDC), 13125 Berlin, Germany

[2] Charité-Universitätsmedizin, 10117 Berlin, Germany

[3] Proteomics Technology Platform, Max-Delbrück-Center for Molecular Medicine and Berlin Institute of Health at Charité Universitätsmedizin Berlin, Berlin, Germany

[4] Department of Rheumatology and Clinical Immunology, Sapporo Medical University School of Medicine, Sapporo, Japan

[5] Department of Pathology, Sapporo City General Hospital, Sapporo, Japan

[6] Biomedical Primate Research Centre (BPRC), Rijswijk, The Netherlands

[7] Department of Gastroenterology and Hepatology, Sapporo Medical University School of Medicine, Sapporo, Japan

[8] Department of Rheumatology and Clinical Immunology, Sapporo City General Hospital, Sapporo, Japan

[9] BIH Center for Regenerative Therapy (BCRT), Berlin Institute of Health @ Charité, Berlin, Germany

[10] Princess Máxima Center for Pediatric Oncology, Heidelberglaan 25, 3584, CS, Utrecht, The Netherlands

[11] Oncode Institute, Utrecht, The Netherlands

[12] DZHK (German Centre for Cardiovascular Research), Partner Site Berlin, 13347 Berlin, Germany

[13] Helmholtz Institute for Translational AngioCardioScience (HI-TAC) of the Max Delbrück Center for Molecular Medicine in the Helmholtz Association (MDC) at Heidelberg University, Heidelberg, Germany

[14] Senior Author

[#] Corresponding author

Correspondence:

1. Masatoshi Kanda, MD, PhD. - (M.K)
2. Norbert Hübner, MD, PhD. - (N.H)

**Table of Contents**

**Supplementary Figures**

**Figure S1:** Ribosome occupancy of *LINC01503* across 4 tissues, related to Figure 1

**Figure S2:** Mass Spectrometry detection of tryptic peptide, related to Figure 1

**Figure S3:** N-terminus FLAG-tagged LINC01503-MP detection, related to Figure 1

**Figure S4:** N/C-terminus FLAG-tagged LINC01503-MP detection, related to Figure 1

**Figure S5:** Co-localisation of overexpressed LINC01503-MP with mitochondria, related to Figure 3

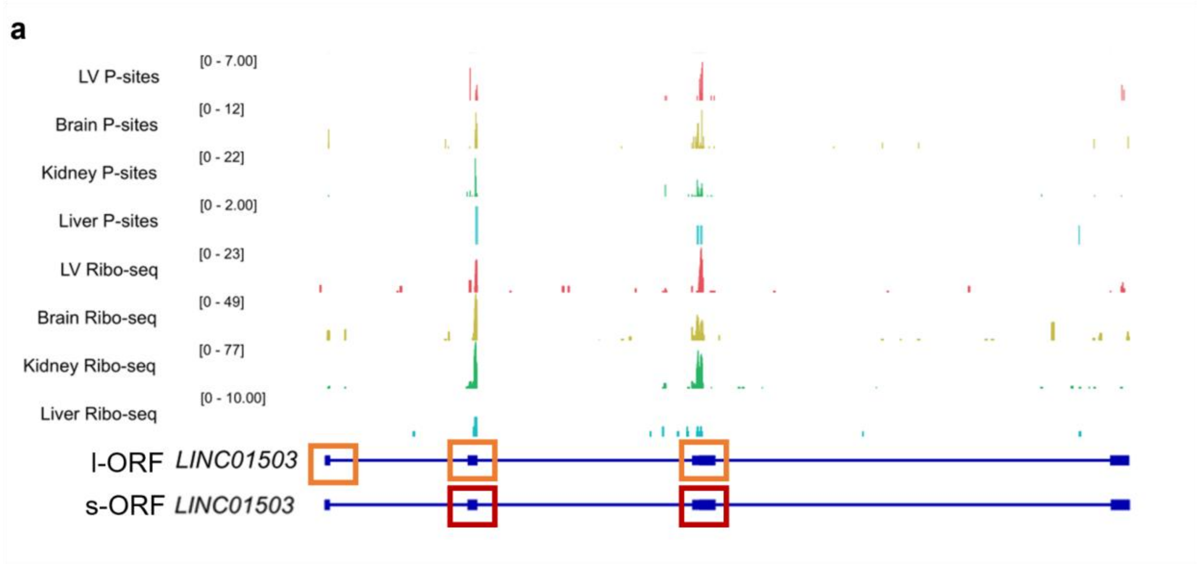


**Figure S1 - Ribosome occupancy of *LINC01503* across 4 tissues, related to Figure 1**

a. Genome coverage tracks visualizing Ribo-seq reads and P-site positions on human *LINC01503* in the four considered tissues. Both ORFs showed high ribosome occupancy in the four tissues evaluated with Ribo-Seq. The number inside of the square bracket indicates the range of counts. Orange boxes indicate the long ORF (l-ORF) encoded by *LINC01503*, spanning three exons. Red boxes indicate the short ORF (S-ORF) encoded by *LINC01503*, spanning two exons.

a

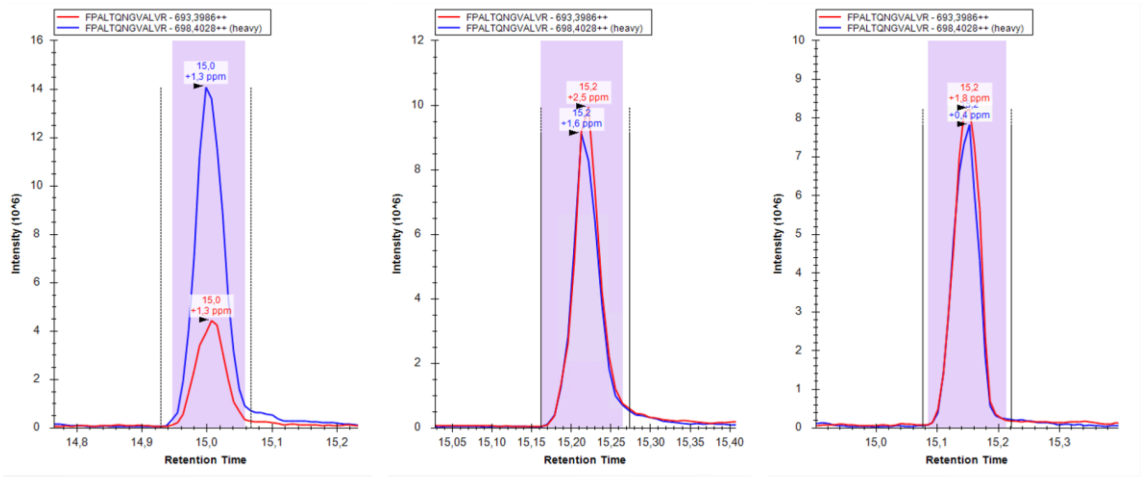


**Figure S2 – Mass Spectrometry detection of tryptic peptide, related to Figure 1**

a. PRM mass spectrometry verification of unique peptide ‘FPALTQNGVALVR’ (heavy) of overexpressed LINC01503-MP in HCT116 cells (n=3).

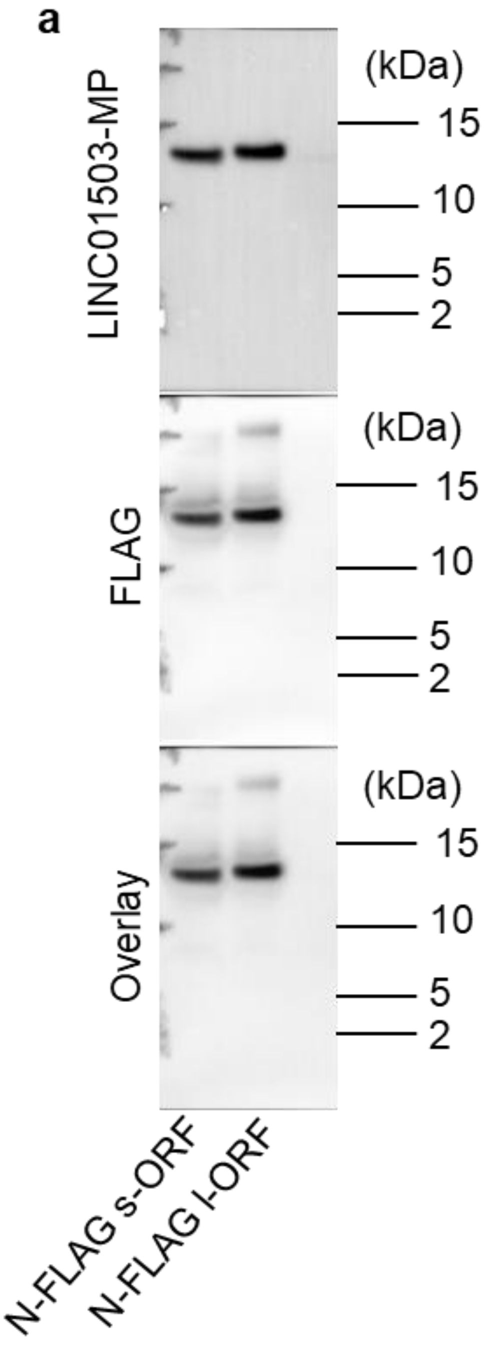


**Figure S3 – N-terminus FLAG-tagged LINC01503-MP detection, related to Figure 1**

b. Western blotting of N-terminus FLAG-tagged LINC01503-MP detected by anti-FLAG Ab and anti-LINC01503-MP Ab. The overlay of both blots showed that both antibodies detected the same fusion microprotein.

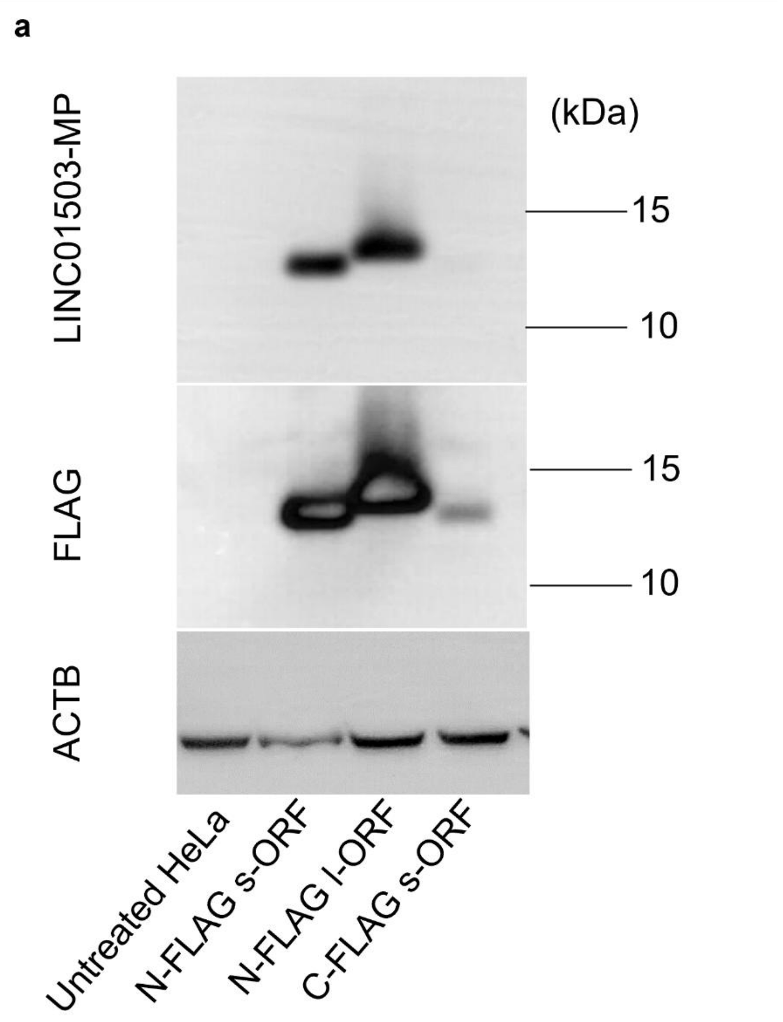


**Figure S4 – N/C-terminus FLAG-tagged LINC01503-MP detection, related to Figure 1**

a. Western blotting of N-terminus and C-terminus FLAG-tagged LINC01503-MP in HeLa cells. Beta-actin (ACTB) is the loading control.

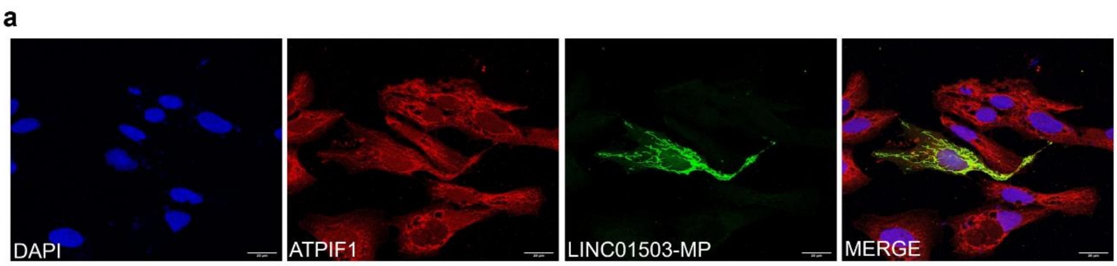


**Figure S5 – Localization of overexpressed LINC01503-MP with mitochondria, related to Figure 3**

a. Immunofluorescence (IF) staining of overexpressed N-terminal FLAG-LINC01503-MP (green) co-localising with ATPIF1 (red) (a mitochondrial protein) in HeLa cells. Scale bar: 20µm.
